# Early Sleep-Dependent Sensory Gating in the Olfactory System

**DOI:** 10.1101/2025.11.04.686664

**Authors:** Diego Serantes, Diego Gallo, Anttonella García, Joaquín González, Mateo Mendoza, Patricia Lagos, Pablo Torterolo, Matías Cavelli

## Abstract

Disconnection from the external world is a defining feature of sleep. Although most models attribute sensory gating to thalamocortical mechanisms, the olfactory system—bypassing the thalamus—offers a unique window into earlier stages of sensory disconnection. Here, we combine chronic and acute recordings in rodents to test whether nasal sensory inputs are internalized during sleep. We show that respiration-locked potentials and gamma activity in the olfactory bulb are strongly modulated by brain state: they diminish during sleep and reappear during wakefulness and cortical activation. This gating occurs independently of respiratory dynamics and arises near the first synapse of the olfactory pathway. Finally, we find that neocortical slow-wave activity correlates with reduced connectiveness, with coupling progressively vanishing as sleep deepens. These findings reveal a sleep-dependent sensory gating mechanism at early stages of a non-thalamic pathway, providing new insights into the neural substrates of sensory disconnection during sleep.

## Introduction

Sleep is a universal state marked by a profound decoupling from the external world. (Andrillon and Kouider, 2020; Cirelli and Tononi, 2024; Coenen, 2024; Velluti, 1997). Despite the high cost of sensory disconnection, the brain willingly enters this state daily—implying that this isolation serves a vital function, such as supporting synaptic homeostasis or offline neural processing (Girardeau and Lopes-Dos-Santos, 2021; Tononi and Cirelli, 2014). However, the precise *when*, *where,* and *how* of sensory gating remain elusive (Cirelli and Tononi, 2024). While most models focus on thalamic and cortical gating, especially during non-REM (NREM) sleep (Andrillon and Kouider, 2020; Coenen, 2024; McCormick and Bal, 1997, 1994a; Olcese et al., 2018), some evidence suggests that earlier, even peripheral stages of sensory disconnection may also occur (Foo and Mason, 2003; Schröder et al., 2020; Velluti, 1997).

The olfactory system, a sensory pathway that lacks a first-order thalamic relay (Kay and Sherman, 2007), provides an ideal model for testing whether sensory gating can occur at the earliest stages of sensory processing (Cirelli and Tononi, 2024; Merrick et al., 2014). Odor information reaches the olfactory bulb (OB) and primary olfactory cortex via only one or two synapses from the external world (Cirelli and Tononi, 2024; Yamaguchi, 2017). This unique anatomy implies that sensory gating may occur at a lower hierarchical level than in other sensory modalities. The OB receives mechanosensory and chemosensory inputs from the nasal epithelium that are tightly coupled to the respiratory cycle (Connelly et al., 2015; González et al., 2023b; Grosmaitre et al., 2007; Iwata et al., 2017), generating a slow, respiration-locked potential and associated gamma activity even in odorless conditions (Adrian, 1942; Bressler and Freeman, 1980; Cavelli et al., 2020; Gonzalez et al., 2024; González et al., 2023b; Iwata et al., 2017; Rojas-Líbano et al., 2014; Tort et al., 2025). This respiration-locked potential has been proven to be essential for olfactory coding and perception (Iwata et al., 2017; Jordan et al., 2018; Kay, 2014; Kepecs et al., 2006; Shusterman et al., 2011; Smear et al., 2011)

The internalization of this respiration-locked potential entrains activity in wide cortical and limbic networks (Ito et al., 2014; Karalis and Sirota, 2022; Tort et al., 2025, 2018; Zelano et al., 2016), and is abolished by blocking nasal airflow (Cavelli et al., 2020; Karalis and Sirota, 2022; Lockmann et al., 2016). Several studies have reported a reduction of this signal during sleep and anesthesia (Cavelli et al., 2020; González et al., 2023a; Homeyer et al., 1995; Jessberger et al., 2016; Khazan et al., 1967), raising the possibility of an early state-dependent modulation of olfactory processing (Tort et al., 2025). On the other hand, Schreck et al. (2022), showed that optogenetic stimulation of olfactory sensory neurons (OSNs) elicited stronger responses in the OB and other brain areas during NREM and REM sleep than during wakefulness. The authors suggest that a centrifugal gating mechanism does not operate during sleep. Instead, the slower and less forceful inhalations typical of sleep reduce odorant access to the epithelium, thereby acting as a partial peripheral gate.

Here, we combine chronic recordings across natural sleep-wake cycles with acute urethane preparations to test whether the respiration-locked activity in the OB persists across brain states and whether its attenuation reflects a genuine gating mechanism or merely changes in nasal airflow. We demonstrate that this coupling is reduced or abolished as sleep deepens and reemerges during wakefulness or urethane-induced activation, even under matched sensory input conditions. These findings support the hypothesis of a functional, early-stage sensory disconnection mechanism operating near the periphery of a non-thalamic sensory pathway.

## Results

### Sleep disrupts respiration-locked potential in the olfactory bulb

We recorded epidural EEG signals from the OB and multiple neocortical areas in freely behaving rats, along with neck electromyogram (nEMG) and a respiratory signal derived from diaphragmatic EMG (dEMG; Fig. S1). Recordings were obtained across spontaneous wakefulness, NREM, and rapid eye movement (REM) sleep (Fig. 1a–b). During wakefulness, the OB displayed a prominent low-frequency potential phase-locked to the respiratory cycle. This signal was strongly attenuated during both NREM and REM sleep (Fig. 1b), reappearing only during brief REM sleep episodes or during microarousals (Fig. S2). Spectral analysis confirmed a robust drop in coherence between OB activity and respiration during sleep (Fig. 1c). As expected, respiratory frequency generally declined and stabilized during sleep (Fig. 1d). Several studies have shown that OB gamma-band activity (40-100 Hz) is modulated by respiration during wakefulness (Adrian, 1942; González et al., 2023b; Tort et al., 2025, 2018).

**Figure 1.**
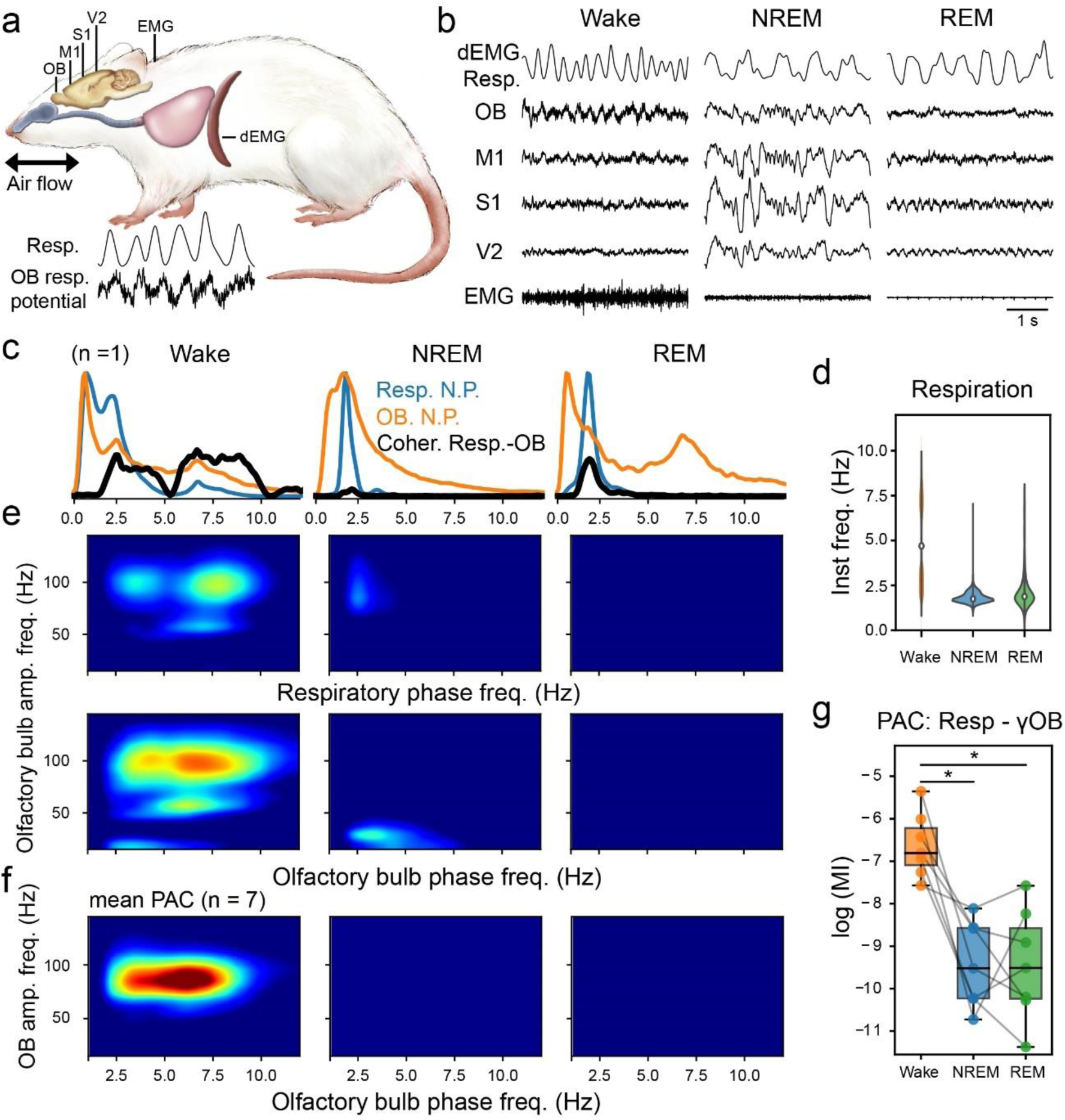
OB respiratory potential during the natural sleep-wake cycle. ***a.*** Schematic representation of electrode placement in the brain and electromyogram recordings from neck muscles (EMG) and diaphragm (dEMG). Bottom: representative example of a respiratory signal and the OB respiratory potential. ***b.*** From top to bottom: recordings of respiratory activity derived from the dEMG, cortical electrical activity, and EMG during Wake, NREM, and REM sleep. Note the presence of respiratory potentials in the OB during wakefulness and their absence during both sleep states. ***c.*** Representative example of normalized power spectra of the OB (orange), respiration (blue), and coherence spectrum between the two signals (black) during wakefulness and sleep. Note the presence of coherence during wakefulness, demonstrating the internalization of the respiratory signal, and its drop during sleep. The x-axis is the same as in panels **e** and **f**. Note that the **c** plots extend to lower frequencies than the panels below, due to their higher frequency resolution. ***d.*** Representative example of instantaneous respiratory frequency across different states of the sleep-wake cycle. ***e.*** Phase-amplitude coupling comodulogram across the sleep-wake cycle for a representative animal. The y-axis shows the amplitude of high-frequency activity. The x-axis shows the phase frequency of the respiratory signal extracted from the dEMG (top panel) and the respiration-locked potential in the OB (bottom panel). Note the amplitude modulation of high-frequency activity during wakefulness, which is lost during sleep. ***f.*** Same as in **e**, but averaged across multiple animals (n = 7). ***g.*** Quantification of respiration-gamma (60-95 Hz) modulation using the modulation index. * p < 0.05. Resp: Respiration. M1: Primary Motor Cortex. S1: Primary Somatosensory Cortex. V2: Secondary Visual Cortex. N.P.: Normalized Power. PAC: Phase–Amplitude Coupling. Amp. freq.: Amplitude Frequency.

Here, we show that this phase-amplitude coupling (PAC) also decreases during both NREM and REM sleep (Fig. 1e–g and S3) (Cavelli et al., 2020; González et al., 2023a). Notably, both coherence and PAC were preserved across a broad range of respiratory frequencies during wakefulness (Fig. 1c and e, Fig. S3). These results demonstrate that sleep disrupts both coherence and respiration–gamma coupling in the olfactory system.

### Urethane anesthesia models NREM-like sensory gating in the olfactory bulb

To complement our findings during natural sleep, we leveraged urethane anesthesia, a preparation that reproduces spontaneous alternations between activated and slow-wave cortical states in the absence of movement or behavioral confounds. This model permits more invasive and controlled recordings, allowing for mechanistic insight into sensory processing across states. We recorded from the same seven rats following intraperitoneal injection of urethane (1.25 g/kg). This preparation produces spontaneous and cyclic alternations between two electrocortical states: an activated state (ASt), characterized by low-amplitude fast activity, and a slow-wave state (SWSt), dominated by high-amplitude slow-wave oscillations (Fig. 2a,b). Historically, this model has been used to approximate the natural sleep cycle, with ASt and SWSt interpreted as REM-like and NREM-like states, respectively (Clement et al., 2008; Pagliardini et al., 2013). Under urethane, we observed a clear respiration-locked potential in the OB during the ASt that was absent during SWSt (Fig. 2a). These changes coincided with spontaneous shifts into a cortical SWSt and were accompanied by fluctuations in respiratory efforts and frequency (Fig. 2b–d), consistent with previous characterizations of urethane-induced state cycling (Pagliardini et al., 2013). Spectral analyses confirm a decrease in the OB respiration-locked potential during SWSt, resulting from a significant reduction in coherence between the OB and the respiratory signal (Fig. 2f,g). Moreover, PAC between respiration and OB gamma amplitude was present during ASt but disappeared during SWSt (Fig. 2h,i). These findings show that urethane reproduces the state-dependent gating observed between wake and NREM sleep, but not the REM-related decoupling from respiration (Cavelli et al., 2020; González et al., 2023a), making it a helpful model for gating, but not for sleep cycling studies, at least at the olfactory system level.

**Figure 2.**
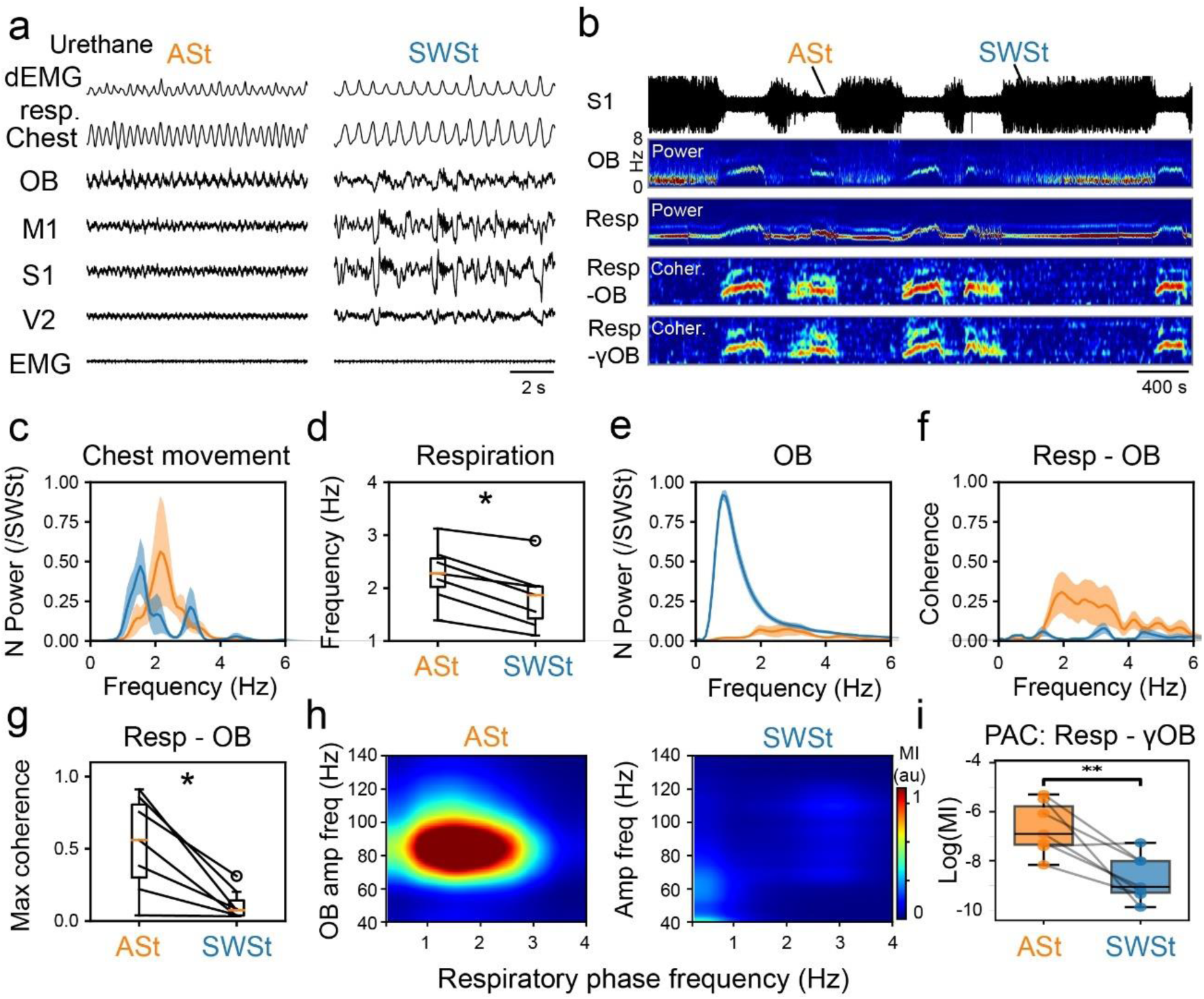
Internalization of the respiratory rhythm during the urethane-activated state and its loss during the slow-wave state. ***a.*** Electrophysiological patterns induced by urethane administration. From top to bottom: respiratory activity extracted from the diaphragmatic EMG (dEMG), chest movement recorded with a motion transducer, cortical electrical activity, and neck EMG. Electrocorticographic traces clearly show an activated state (ASt) with low amplitude and high frequency resembling wakefulness, and a slow-wave state (SWSt) with high amplitude and low frequency resembling NREM sleep. Note the presence of respiratory potentials in the olfactory bulb (OB) during ASt and their absence during SWSt. ***b.*** From top to bottom: *1* – Compressed S1 cortical recording. Note that amplitude fluctuations in the raw trace distinguish the two urethane-induced states. *2* – Power of the olfactory bulb. *3* – Power of diaphragmatic respiration. Note that changes in respiration are coupled with fluctuations in OB power. *4* – Coherence between respiration and OB. *5* – Coherence between respiration and gamma-band amplitude (60-95 Hz) in the OB (Cross-frequency coupling). Note that gamma modulation occurs during ASt but not during SWSt. ***c.*** Normalized power spectrum (mean ± std) of respiration in both states. The normalization was made to the maximum value of SWSt (/SWSt). ***d.*** Instantaneous respiratory frequency averaged across states (n = 7). * p < 0.05. ***e.*** Normalized power spectrum of the OB. ***f.*** Spectral coherence between respiration and OB. ***g.*** Maximum coherence values between respiration and OB EEG, across states. Note the significantly higher coherence during ASt, which decreases during SWSt, resembling the pattern observed during wakefulness and sleep. ***h.*** Phase–amplitude coupling (PAC) comodulogram for both states. The y-axis shows the amplitude of high-frequency activity; the x-axis shows the phase frequency of respiration. Note the strong coupling in ASt and its absence in SWSt. ***i.*** Modulation index quantification. Note the significant differences between the two states. ** p < 0.001 Resp: Respiration. M1: Primary Motor Cortex. S1: Primary Somatosensory Cortex. V2: Secondary Visual Cortex. N Power: Normalized Power.

### The recorded gated signal originates from the OB local circuitry

To confirm that the gated respiratory potential recorded with surface electrodes over the OB reflects local activity rather than volume-conducted signals, we performed acute recordings using linear silicon probes in urethane-anesthetized rats. Respiration was measured using both a nasal thermistor and a piezoelectric chest sensor to assess respiration-locked potentials in the olfactory bulb. LFP inversion and CSD analyses along the dorsoventral axis revealed clear respiration-locked sinks and sources during the ASt, which disappeared during the SWSt, confirming the local origin of the signal (Fig. 3a-b). LFPs recorded across OB layers showed strong respiratory modulation during ASt that disappeared in SWSt (Fig. 3a-c). Coherence between respiration and OB activity, computed from both nasal airflow (thermistor) and thoracic movement (chest piezo sensor), was high during ASt and significantly reduced in SWSt (Fig. 3d). PAC between respiratory phase and OB high-frequency activity followed the same pattern, with robust modulation during ASt and a marked drop in SWSt (Fig. 3e,f). Notably, the results were consistent across respiratory signals (nasal thermistor or thoracic movement) and neural recording methods (epidural EEG or depth-resolved LFP/CSD). These results confirm that the respiration-locked potential and gamma activity in the OB are locally generated and strongly modulated by brain state.

**Figure 3.**
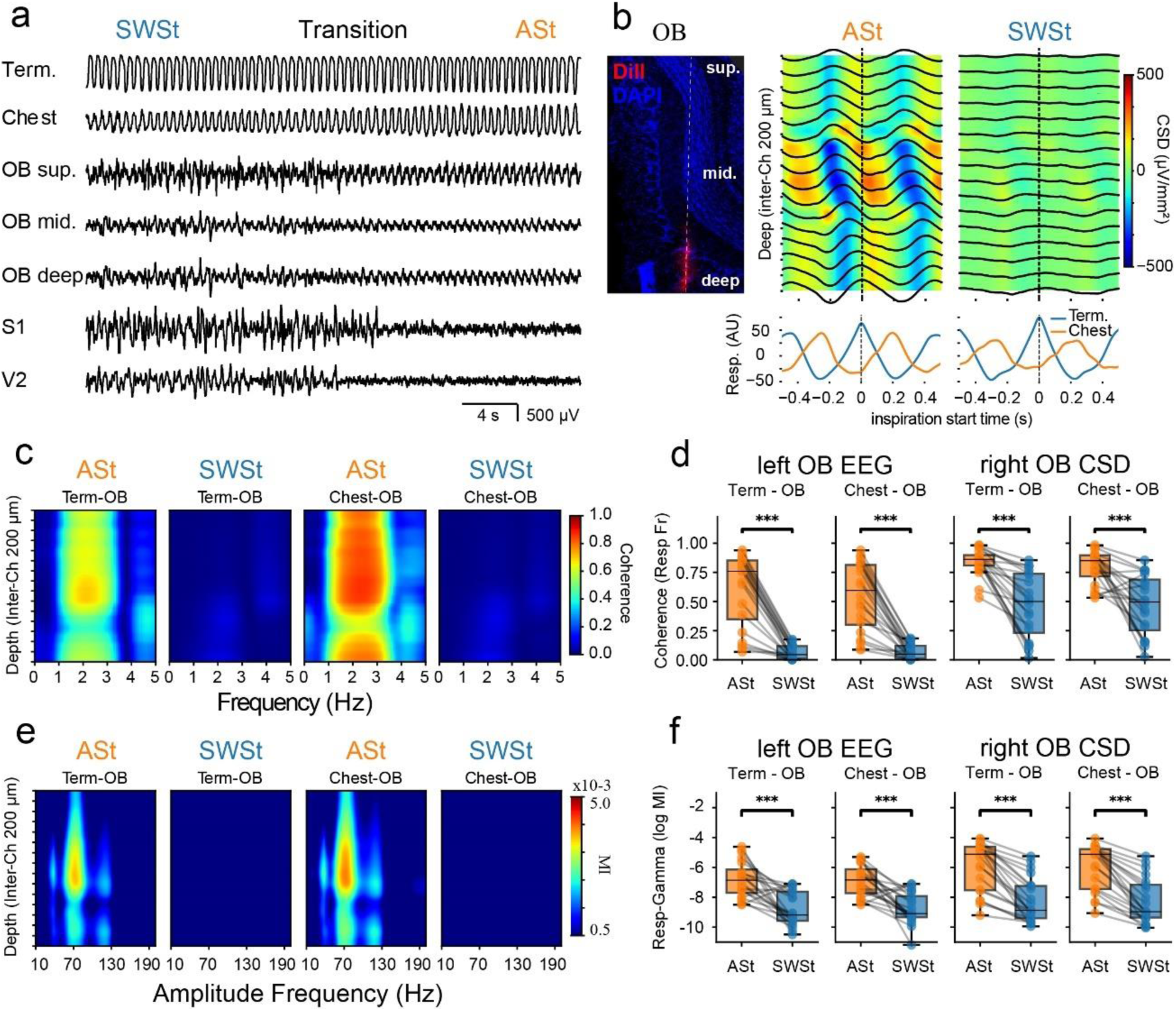
Respiration-locked potential and gamma activity are generated locally in the olfactory bulb. ***a.*** Representative recording showing a transition from a slow-wave state (SWSt) to an activated state (ASt). From top to bottom: nasal airflow recorded with a thermistor (Term.), thoracic wall movement (Chest), local field potentials (LFPs) from superficial, middle, and deep layers of the OB recorded with a Neuronexus probe, and cortical activity from S1 and V2 used to determine brain states. ***b.*** Left: Coronal section of the olfactory bulb showing the trajectory of a DiI-labeled Neuronexus probe (red) and DAPI-stained nuclei (blue). Center and right: Average current source density (CSD) and LFP (black traces) aligned to the onset of nasal inspiration (dashed line) during ASt and SWSt, respectively. Prominent respiration-locked oscillations with alternating current sinks and sources are observed during ASt, while such modulation is nearly absent during SWSt. Bottom: Mean respiratory signals from the nasal thermistor (blue) and thoracic sensor (orange), aligned to inspiration onset. ***c.*** Spectral coherence maps between the respiratory signals (thermistor and chest) and OB LFPs across depth, during ASt and SWSt. Coherence is high during ASt and markedly reduced during SWSt. The same experiment as **b**. ***d.*** Group-level coherence between respiration (thermistor or chest) and either left OB epidural EEG or right OB CSD. For this analysis, the CSD was constructed using the three most distant and equally spaced channels. Coherence was significantly higher during ASt than SWSt for both respiratory signals and recording types. ***e.*** Phase–amplitude comodulograms showing the coupling between respiratory phase (thermistor or chest) and high frequency amplitude across OB layers. Same experiment that **b**. ***f.*** Modulation index (MI) quantifying phase–amplitude coupling between respiration and gamma-band (60–90 Hz) activity, computed from left OB EEG and right OB CSD. For this analysis, the CSD was constructed using the three most distant and equally spaced channels. Significant reductions in MI were observed during SWSt compared with ASt across both respiratory sources and recording types. *** p < 0.0001. Inter-Ch: Inter-Chanel. sup., superficial; mid., middle.

### Nasal airflow is necessary but not sufficient to generate the olfactory bulb respiratory potential

Previously, we confirmed that both thoracic and nasal respiratory signals reliably capture the respiration-locked potential in OB (Fig. 3). To directly test whether nasal airflow alone is sufficient to drive the respiratory potential in the OB, we used a double-tracheotomy preparation that dissociates thoracic breathing from nasal airflow (Fig. 4a–b). While animals continued to breathe spontaneously through the lower tracheal cannula (open to the atmosphere), an artificial sniffing system delivered suction airflow pulses through the upper cannula into the nasal cavity, mimicking natural inhalations independently of thoracic movement. LFPs were recorded across OB layers using a laminar probe, alongside epidural EEG (Fig. 4c–e). During ASt, OB LFPs exhibited robust oscillations phase-locked to the artificial nasal airflow across most tested stimulation frequencies (Fig. S4). This stimulation-locked signal vanished when the stimulus was turned off (Fig. 4c,e; Fig. S4), and decreased upon transition to SWSt, despite constant inspiratory efforts across states (Fig. 4d). Spectral coherence between OB activity and the nasal airflow signal was high during ASt and decreased during SWSt. In contrast, coherence with the thoracic signal was generally low across states (Fig. 4f). Phase–amplitude coupling analyses mirrored this selectivity: OB gamma activity was modulated by nasal airflow only during ASt (Fig. 4f). Occasionally, a weak OB–chest coherence was observed in a small subset of experiments, but it lacked a corresponding chest–gamma coherence and was not consistent across experiments (Fig. 4h). These results show that nasal airflow is required but not sufficient: OB responses depend on brain state and vanish during SWSt, even when airflow parameters are kept constant.

**Figure 4.**
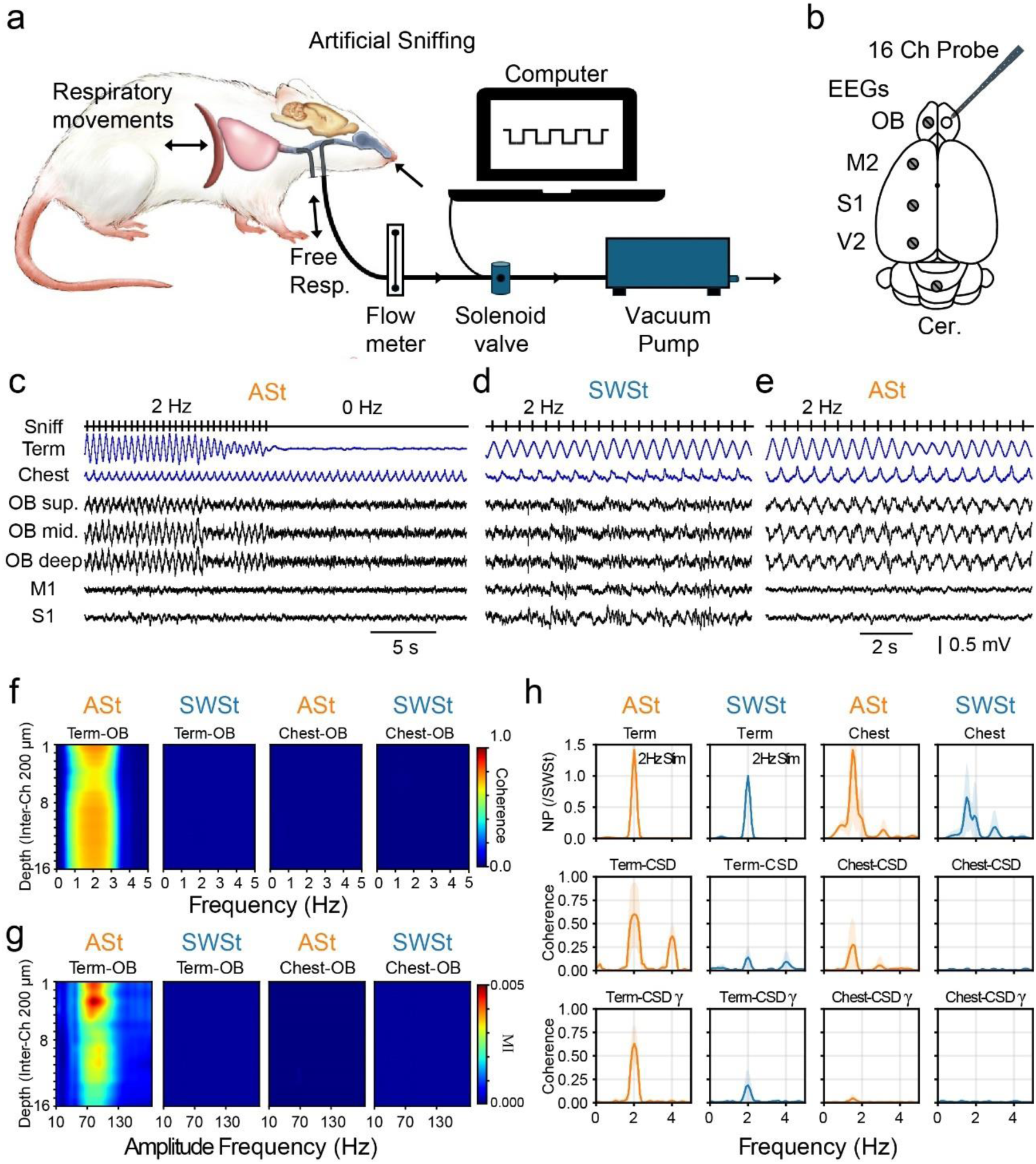
The internalization of respiration-locked potential in the OB requires both nasal airflow and cortical activation. ***a.*** Schematic of the double tracheotomy preparation. The lower tracheal cannula was left open to the environment, allowing spontaneous lung ventilation. In contrast, the upper tracheal cannula was connected to an artificial sniffing system that delivered controlled nasal airflow independent of thoracic movement. ***b.*** Schematic dorsal view of the rat brain illustrating the anteroposterior distribution of epidural electrodes and LFP probe. primary motor cortex (M1), primary somatosensory cortex (S1), and secondary visual cortex (V2). In the right hemisphere, a 16-channel Neuronexus probe was inserted into the OB for laminar LFP recordings. ***c.*** Representative recording during an activated state (ASt). From top to bottom: cortical EEG from M1 and S1; four LFP channels from the OB; TTL signal from the microcontroller triggering the solenoid valve (Pump); thermistor signal (Term) monitoring nasal airflow; and chest movement signal (Chest) reflecting spontaneous respiration. Note that stimulation at 2 Hz during the first half of the recording evoked a stimulus-locked potential, which vanished upon deactivation. **d-e.** Nasal airflow remained constant throughout the recording, while the cortical state spontaneously transitioned from SWSt to ASt. Note the emergence of respiration-locked OB oscillations only during ASt. ***f.*** Coherence between OB LFPs and both respiratory signals (thermistor and chest) across depth. During ASt, nasal airflow (Term) was coherent with OB activity at the stimulation frequency, whereas chest movement showed no coherence in either state. ***g.*** Modulation index (MI) between respiratory phase and OB gamma amplitude (60–90 Hz). Robust phase–amplitude coupling was observed exclusively between nasal airflow and olfactory bulb gamma activity at the sniffing frequency, and only during the activated state (ASt). ***h.*** Population summaries (mean ± s.d.) of spectra in 0–5 Hz. Top row: normalized power of the respiratory signals (Term/Chest), normalized within each animal to the SWSt maximum for that signal. Middle row: coherence between each respiratory signal and the OB CSD. Bottom row: coherence between each respiratory signal and the OB gamma envelope (60–90 Hz). sup., superficial; mid., middle.

### Cortical slow-wave activity predicts the strength of the respiration-locked potential in the olfactory bulb

To determine whether sensory gating in the olfactory system scales with sleep depth, we computed the coherence between nasal respiration and the LFP in the OB across the entire recording. We also calculated the modulation index (MI) between nasal respiration and the gamma amplitude. Both analyses were conducted in 5-second non-overlapping windows, each classified as ASt, Transitional (Trans), or SWSt. To measure slow-wave activity (a proxy for sleep depth), delta power (0.5–4 Hz) was extracted from each window (Cavelli et al., 2023). To determine whether the coupling between the respiration and OB signals was statistically significant, we performed a surrogate-data analysis by generating phase-randomized signals and comparing the observed coherence and MI with the surrogate distribution.

In all experiments, we found a robust negative correlation between delta power and respiration-LFP coherence (Fig. 5a), indicating that as slow-wave activity increases, OB activity becomes less connected to the outside world. A similar pattern was observed for the MI between respiration and gamma amplitude (Fig. 5a), further supporting a graded attenuation of respiratory drive onto olfactory circuits with increasing slow-wave activity. It is noteworthy that this coherence and MI values fell below significance mostly during SWSt (Fig. 5b). Specifically, only 9.23 ± 6.72% (mean ± std) of the SWSt windows reached significance for coherence (vs. 26.05 ± 12.92% in ASt and 20.41±15.29% in Trans). Only 9.64 ± 9.49% reached significance for MI (vs. 33.72 ± 21.18% in ASt and 15.19 ± 16.77% in Trans).

**Figure 5.**
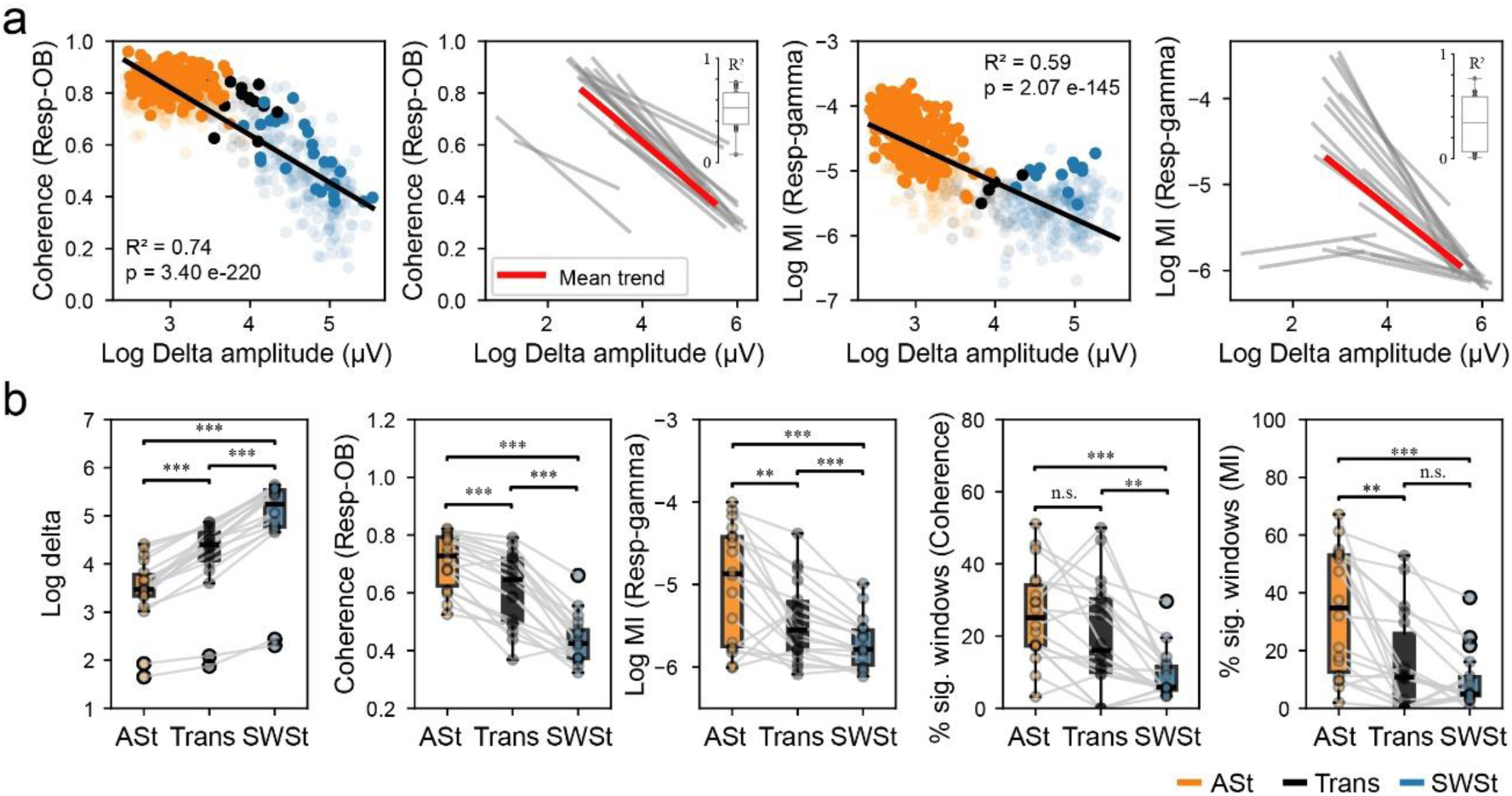
Delta power determines internalization of the respiratory cycle. ***a.*** Left: Example from a single animal showing the negative correlation between delta power and nasal respiration – OB LFP coherence. Orange, black, and blue circles represent ASt, transition, and SWSt epochs, respectively; color-transparent dots show non-significant windows (surrogate analysis). Second panel: Significant linear fits (p< 0.001) of this relationship across all animals and experiments, with the mean trend highlighted in red and the distribution of R² values shown in the inset. Third panel: Example from a single animal showing the relationship between delta power and the modulation index (MI; OB gamma–respiration coupling). Rightmost panel: Same relationship across all animals and experiments, with mean trend and R² distribution inset. **b** Group-level comparisons of delta power, respiration–OB coherence, and OB gamma–respiration MI across ASt, transition, and SWSt epochs. The last two panels display the percentage of significant windows for the delta–coherence and delta–MI relationships, respectively. Lines link data from the same animal. Statistical significance: **p < 0.01; ***p < 0.001; n.s., not significant.

## Discussion

### Summary of the main findings

Our results demonstrate a brain-state-dependent attenuation of the respiration-locked activity in the OB, a signal that reflects mechanosensory and chemosensory inputs from the nasal epithelium. This attenuation occurs both during natural sleep and in the urethane-induced slow-wave state and persists even under matched sensory inputs. We demonstrate that this gating effect is expressed locally within the OB (without volume conduction), requires nasal airflow, is independent of respiratory frequency, and scales negatively with slow-wave activity, a proxy for sleep depth. Notably, the coupling between respiration and OB activity, measured by both coherence and MI, frequently falls below chance levels during deep slow-wave states, indicating a functional disconnection from the outside world.

### Peripheral vs. central gating in the olfactory system

Although not all models agree on the precise locus of sensory gating, the prevailing view of sensory disconnection during sleep emphasizes thalamocortical gating mechanisms, wherein the sleep spindle activity and/or the slow wave-related bistability of thalamic and cortical neurons play a central role in interrupting the causal chain triggered by externally and internally generated inputs (Andrillon and Kouider, 2020; Cavelli et al., 2023; Fernandez and Lüthi, 2020; Koch et al., 2016; McCormick and Bal, 1997, 1994b; Olcese et al., 2018; Pigorini et al., 2015; Siclari et al., 2017; Steriade et al., 1993; Tononi et al., 2016). However, the olfactory system bypasses the thalamus in its primary ascending pathway (Cirelli and Tononi, 2024; Yamaguchi, 2017), raising the question of whether gating can occur at earlier stages of processing. Our findings suggest that gating occurs at or before the OB, a site only one synapse away from the nasal epithelium. These results align with early work that describes the reduction of OB potentials during sleep (Cavelli et al., 2020; González et al., 2023a; Jessberger et al., 2016; Khazan et al., 1967), which was not always interpreted as active gating or sensory disconnection.

Recent evidence further complicates the current understanding. Schreck et al. (2022) showed that random optogenetic activation of olfactory sensory neurons (OSNs) produces robust activation of the OB and downstream cortical targets during sleep, including stronger responses in NREM than in wake. They proposed that slower, less forceful inhalations during sleep might reduce epithelial sampling, constituting a passive peripheral gate. Our data, by contrast, reveals a gating mechanism that persists even under artificial, matched sniffing conditions, arguing for an active, state-dependent gating process localized in the OB or earlier in the olfactory pathway.

### Urethane as a model for testing sensory gating

Urethane anesthesia induces spontaneous alternations between activated and slow-wave-like cortical states, which have been classically interpreted as analogs of REM and NREM sleep, respectively (Clement et al., 2008; Pagliardini et al., 2013; Ward-Flanagan et al., 2024). However, this interpretation has been increasingly challenged (Brankačk et al., 2025; Mondino et al., 2022). This (Fig. 1 and S2) and our previous studies (Cavelli et al., 2020; González et al., 2023a) have shown that the respiration-locked potential and the phase-amplitude coupling between the respiratory phase and gamma oscillations are strongly attenuated during natural REM sleep (Fig. 1 and S2), and even during REM sleep induced by carbachol injection in the Nucleus Pontis oralis (NPO) (Cavelli et al., 2020). These findings suggest that there is no functional equivalence between urethane-activated states and REM sleep, at least from the perspective of olfactory sensory processing.

We show that, under urethane anesthesia, the respiration-locked potential is suppressed during the slow-wave state and re-emerges during cortical activation, despite matched peripheral input. This pattern mirrors what is observed between NREM sleep and wakefulness but contrasts with the dissociation typically seen in natural REM sleep. Thus, we argue that urethane cycling does not recapitulate the sensory dynamics of the natural sleep cycle and should not be used to model REM-state olfactory processing.

Nevertheless, urethane anesthesia provides a powerful model for isolating the mechanisms of sensory gating. Its stability and lack of behavioral confounds allow for invasive manipulations — such as tracheotomy, artificial sniffing, and laminar recordings — that would be difficult to implement across natural sleep–wake cycles. From this perspective, urethane preparations provide an ideal framework for studying state-dependent gating mechanisms under tightly controlled sensory conditions.

### Sensory disconnection in the olfactory system: Revisiting evidence

Behavioral studies have consistently shown that controlled olfactory stimuli are less likely to elicit arousal during sleep, anesthesia, or coma, especially when presented in isolation (Badia et al., 1990; Carskadon and Herz, 2004; Mao et al., 2024; Tseng et al., 2022). At the same time, multiple studies demonstrate that olfactory cues delivered during sleep can enhance memory consolidation and modulate electrophysiological sleep markers (Gaeta and Wilson, 2022; Rasch et al., 2007; Rihm et al., 2014). On the other hand, recent work reports robust arousals to certain strong olfactory stimuli (e.g., alcohol), although contextual or multimodal factors may contribute (Mao et al., 2024). In this sense, we found that ∼9% of SWSt windows still show some degree of connectivity with the outside world.

Electrophysiological work in both natural sleep and anesthesia has revealed state-dependent reductions in odor-evoked responses and respiratory entrainment (Cavelli et al., 2020; González et al., 2023a; Jessberger et al., 2016; Khazan et al., 1967; Murakami et al., 2005; Tsuno et al., 2008). However, these reductions were rarely interpreted as evidence of sensory disconnection or functional gating. Here, we bridge that gap. Our analyses of coherence and MI between respiration and OB activity across sleep–wake states reveal a quantitative, graded reduction in sensory coupling with increasing slow wave activity. Many windows during SWSt show coherence and MI values that fall below surrogate thresholds, suggesting a functional loss of sensory entrainment. This dynamic gating, tightly linked to cortical slow-wave activity, supports the idea of a graded disconnection process.

Notably, previous work has shown that respiratory inputs—particularly gamma oscillations modulated by the sniff cycle—enhance odor coding and facilitate information transfer between the OB and piriform cortex during wakefulness (Gonzalez et al., 2024; González et al., 2023b). This establishes a direct functional role for the respiratory-locked signals measured in the OB. Therefore, the state-dependent suppression of these signals during slow-wave states likely reflects not only passive decoupling from peripheral input but also an active, functionally meaningful gating mechanism that modulates olfactory processing and sensory awareness during sleep.

### Where is the gate? Remaining uncertainties

While our data highlight the OB as a critical node for state-dependent gating, the precise mechanisms remain unresolved. One plausible source is centrifugal modulation from arousal-promoting systems in the basal forebrain and brainstem, such as the locus coeruleus and raphe nuclei, which project extensively to the OB and are strongly state-dependent in their firing (Aston-Jones et al., 2000; McLean and Shipley, 1987; Vanini and Torterolo, 2021). These neuromodulatory systems may implement centrifugal gain control during sleep-wake transitions, regulating OB responsiveness in a state-dependent manner.

Noradrenaline and serotonin, in particular, have been shown to modulate odor coding at the OB through cell-type-specific and layer-specific mechanisms. Noradrenaline alters mitral cell excitability and odor detection thresholds (Escanilla et al., 2010; Linster et al., 2011), while serotonergic input suppresses transmission from OSNs to mitral cells via local inhibitory circuits (Petzold et al., 2009). Interestingly, this inhibition is input-specific and may selectively attenuate odor-evoked activity. Although monoaminergic systems are traditionally viewed as quiescent during NREM sleep, recent fiber photometry experiments have shown fluctuations in neuromodulatory tone even during deep sleep (Osorio-Forero et al., 2025), indicating that, if they are responsible, their role in OB gating may be more dynamic than previously thought.

Furthermore, anatomical studies demonstrate that centrifugal inputs to the OB are heterogeneous and spatially organized, targeting distinct regions of the glomerular layer (Gómez et al., 2005). This raises the possibility that centrifugal gating is not uniform but spatially selective, influencing specific odor channels depending on behavioral context or state. In addition, a dopaminergic component may contribute to gating. For instance, studies have described a nigro-olfactory projection (Höglinger et al., 2015) and functional impacts of dopaminergic lesions on odor discrimination and respiration-locked signals (Cavelli et al., 2019; Yuan et al., 2023), suggesting that dopaminergic modulation might influence OB sensitivity or gating under certain conditions.

Beyond changes in excitability or gain at centrifugal connections, it is possible—by analogy with thalamic and cortical systems—that the withdrawal of ascending arousal systems during sleep (Vanini and Torterolo, 2021), and the resulting spindles and bistability (Amzica and Steriade, 1998; Steriade et al., 2001, 1993), may underlie the observed gating (Murakami et al., 2005). Although this remains speculative in the olfactory system, it is conceivable that local sleep patterns themselves could prevent the propagation of sensory-evoked activity, effectively disrupting causal chains triggered by sensory inputs (Cavelli et al., 2023; Massimini et al., 2005; Matosevich and Nir, 2021; Pigorini et al., 2015).

On the other hand, a peripheral origin of gating remains possible (Homeyer et al., 1995; Schreck et al., 2022). Olfactory sensory neurons are subject to autonomic regulation, and the nasal mucosa exhibits rich sympathetic and parasympathetic innervation that could alter epithelial excitability (Baraniuk and Merck, 2009). Changes in vascular tone, mucus viscosity, or receptor conductance could modulate signal transmission before reaching the OB. Neuromodulatory influences at the epithelium have also been documented (Lizbinski and Dacks, 2018; Ma and Luo, 2012), although their behavioral relevance during sleep remains unexplored.

### Potential contribution of internally generated signals

Although the respiration-locked potential in the OB depends primarily on nasal airflow (Fig. 4), it remains plausible that a fraction of this signal arises from internally generated sources, such as corollary discharges or interoceptive inputs linked to respiration (Crapse and Sommer, 2008; Li and Xu, 2022). Supporting this possibility, previous studies have reported residual respiration-locked activity in the olfactory system after tracheotomy (Murakami et al., 2005), and similar residual signals have been observed in other brain regions following epithelial ablation (Karalis and Sirota, 2022).

In our recordings, we occasionally observed low but detectable coherence between thoracic signals and OB CSD activity (Fig. 4h), suggestive of a central contribution. However, our dataset is insufficient to isolate or characterize these internally generated components, and further work will be needed to clarify whether central respiratory signals contribute to olfactory entrainment and how they might vary across sleep–wake states. Such signals, if confirmed, could reflect interoceptive or predictive representations of the respiratory phase, potentially aligning olfactory processing with internally generated expectations during wakefulness (Friston and Kiebel, 2009; Lyons and Gottfried, 2025; Zelano et al., 2011).

### Future directions

Our findings uncover a sleep-depth-dependent gating mechanism operating at or near the entry point of the olfactory system, bypassing the thalamus. A key next step is to pinpoint its origin—whether in the olfactory bulb, the sensory epithelium, or structures such as the nasal mucosa or vascular interfaces. Dissecting the relative contributions of peripheral responsiveness or centrifugal modulation will be essential. Equally important is determining whether this gating merely suppresses the presence of afferent signals or also disrupts their functional role. The disappearance of these signals during slow-wave states may reflect not just passive decoupling but a functional gating process that impairs sensory encoding. Furthermore, analyzing sensory gating during REM sleep —a cortical-activated state characterized by suppression of the respiratory signal in the OB —is also warranted. Future work combining physiological manipulations with behavioral or decoding-based assessments across brain states could test whether gating alters the structure, fidelity, or transmission of odor information. This line of inquiry will help clarify whether sleep-related gating in the olfactory system protects sleep from sensory intrusion or whether it actively shapes sensory processing and memory consolidation during offline brain states.

## Methods

### Experimental animals

Thirty adult Sprague-Dawley rats (320–400 g) were used in this study. The animals were obtained from URBE (Unidad de Reactivos Biológicos para Experimentación), of the School of Medicine, Universidad de la República, Uruguay. All experimental procedures were conducted in agreement with the National Animal Care Law (#18611) and with the ‘Guide to the care and use of laboratory animals’ (8th edition, National Academy Press, Washington D. C., 2010). Furthermore, the Institutional Animal Care Committee approved the animal experimental procedures (N°: 070151-000036-23). Adequate measures were taken to minimize pain, discomfort, or stress of the animals, and efforts were made to use the smallest number of animals necessary to obtain reliable data.

### Surgical procedure

#### Chronic recordings

We employed similar surgical procedures as in our previous studies (Cavelli et al., 2019, 2018; Serantes et al., 2024). The animals were chronically implanted with electrodes to monitor their sleep states and wakefulness. Stereotactic implantation was performed under isoflurane anesthesia (3% induction, 1.5–2.5% maintenance). Using sterile techniques, a midline incision was made to expose the skull. After cleaning the surface with bonding agent (OptiBond), several small burr holes were created in the skull using a dental drill. To record the EEG, stainless steel screw electrodes (0.8 mm in diameter) were screwed into craniotomies, with their tips contacting the brain’s surface (above the dura mater) in different cortical areas and the OB. Six electrodes were located on the neocortex. The electrodes were in the primary motor cortex (M1, L: 2.5 mm, AP: +2.5 mm), the primary somatosensory cortex (S1, L: 2.5 mm, AP: 2.5 mm), and the secondary visual cortex (V2, L: 2.5 mm, AP: 7.5 mm). Another electrode was located over the right OB (L: +1.25 mm, AP: +8-8.5 mm). The last screw was positioned over the cerebellum as an electrode reference. To record the electromyogram (EMG), two electrodes (stainless steel multifilament cable) were inserted into the muscle of the neck. To record respiratory activity, bipolar electrodes were inserted into the diaphragms of two animals. These EMG electrodes were constructed following the design of Shafford et al. and Rojas-Libano et al., who specifically designed them for diaphragm EMG recordings (Rojas-Líbano et al., 2014; Shafford et al., 2006). Briefly, the EMG recording electrode was constructed using a 250 μm diameter stainless steel wire, which was formed into a coil approximately 1–2 mm in length. The coil was then connected to a multi-wire flexible cable and transferred subcutaneously to the skull. The electrodes were soldered into a socket and fixed onto the skull with acrylic cement (Flow). At the end of the surgical procedures, an analgesic (Ketoprofen, 1 mg/Kg s.c.) was administered. Incision margins were kept clean, and a topical antibiotic was applied daily. After the animals recovered from the surgical procedures, they were adapted to the recording chamber for 1 week.

#### Acute recordings

The animals were chronically implanted with electrodes to monitor their electrographic state during acute experiments. Stereotactic implantation was performed under isoflurane anesthesia (3% induction, 1.5–2.5% maintenance). To record the electrographic state, stainless steel screw electrodes were screwed onto the craniotomies in different cortical areas and the OB. Electrodes were positioned identically to those in the chronic animal groups, albeit limited to the left hemisphere. Two references and ground electrodes were placed over the cerebellum (Cer, L: ±1 mm, AP: −12.5 mm). Finally, a small stereotaxic mark accompanied by an acrylic-free perimeter was made on the right olfactory bulb (L: +1 mm, AP: +8.5 mm). This corresponds to the insertion area of the recording probe during the urethane acute experiment. All electrodes were soldered into a socket and secured onto the skull with acrylic cement. Two stainless steel horizontal bars, which serve as stereotactic anchors for future acute experiments, were also secured to the skull with acrylic cement. An analgesic (Ketoprofen,1 mg/Kg s.c.) was administered at the end of the surgical procedure. Incision margins were kept clean, and a topical antibiotic was applied daily. After surgery, the animals recovered for at least one week before acute experiments.

### Experimental sessions

#### Chronic sessions

The animals were housed in acrylic cages lined with wood shavings and placed inside a sound-attenuating chamber that also functioned as a Faraday box. The housing environment was maintained in a temperature-controlled room (21–24°C), with food and water provided *ad libitum*. The experiments were conducted under the light period (9:00 a.m. to 9:00 p.m.) on a 12-h light/dark cycle (lights on at 9 a.m.). All recordings were performed using a rotating commutator, which allowed the rats to move freely within the recording box. Bioelectric signals were amplified (x1000; Grass Model 12), filtered (0.3–300 Hz), sampled (1000 Hz, 16 bits; NI PCI-6220), and stored in a PC using the OpenEphys GUI (Siegle et al., 2017). After several days of baseline recordings, the animals were anesthetized with urethane (1.25 g/kg, i.p.) and placed on a heat mattress therapy pump (Gaymar, T/Pump) with a piezo sensor taped to the chest to record respiratory chest movements and/or a thermistor in front of the nostril.

#### Acute sessions

Following at least one week of recovery, the animals were anesthetized with urethane (1.3-1.5 g/kg, i.p.), put on a heat mattress therapy pump (Gaymar, T/Pump), and positioned in a stereotaxic frame using the previously implanted fixation bars. Small burr holes were drilled over the olfactory bulb (OB) using a dental drill and guided by the reference mark created during the initial stereotaxic surgery. The hole was protected with petroleum jelly (Vaseline). A lineal channel probe (NeuroNexus Technologies; A1×16 10mm-200, A1×32 10mm-100) was inserted perpendicular to the OB surface and used as recording electrodes. Before insertion, the shanks of the probes were coated with a red fluorescent cell-labeling solution (CM-Dil, Thermo Fisher Scientific) for later localization of the electrode tracks in post-mortem histology. After probe alignment, insertion was performed using a micromanipulator until it reached the ventral part of the OB. At the end of the insertion, 20 minutes were used to prepare the recording system and as a tissue stabilization period. Bioelectric signals (EEGs, LFPs, and Units) were amplified (x100), filtered (0.1–7500 Hz), and sampled (30000 Hz, 16 bits) using two 32/16-channel Intan headstages (RHD 16-channel headstage) connected to an OpenEphys acquisition board and stored on PC using OpenEphys GUI (Siegle et al., 2017). EEG arrays were attached to the headstage using an electrode adapter board (Intan 18-pin electrode adapter board). A piezo sensor was taped to the animal to record the respiratory chest movements. A thermistor was positioned in front of the nostril, and a homemade circuit powered it. Bout respiratory signals were amplified (x100) and later entered the OpenEphys acquisition board through the 8-channel ADC board (Siegle et al., 2017). During this recording session, a continuous flow of deodorized air (activated charcoal, 1 L/min) was directed in front of the animal’s nose (Iwata et al., 2017).

#### Tracheotomy and artificial sniffing

In some acute sessions, before positioning the animal in the stereotaxic frame, a double tracheotomy was performed, and two silicone tubes (inner diameter 0.5 mm) were inserted and fixed using super glue. The lower tube allowed rats to breathe freely, while the upper tube was positioned at the nasopharynx. The other end of the upper tube was connected to a solenoid valve, a flow meter, and an air suction pump. The nasal airflow rate was 0.4-1 ml/min and frequency 0.5-10 Hz, with negative pressure pulses applied for half the duration of the sniff cycle. The solenoid valves controlling nasal airflow were regulated using relay circuits and an Arduino (Iwata et al., 2017; Oka et al., 2009). This preparation enables us to control the inspiratory airflow during all electrographic states.

### Histology

At the end of the recording session, rats were intracardially perfused with PBS (phosphate buffer solution with heparin 5000 IU/L) and 4% paraformaldehyde (PFA) in PBS for tissue fixation. Brains were then extracted and processed for histology. After fixation, brains were cryoprotected by exposure to increasing concentrations of sucrose in PBS solutions at 4 °C. Brains were then quickly frozen and sliced in coronal sections (40-50 μm thick) with a cryostat. Sections were mounted with medium containing DAPI (SouthernBiotech; DAPI-Fluoromount-G). To verify the probe location, sections were imaged with an upright epifluorescence microscope (Leica; DM2500) (Cavelli et al., 2023).

### Data analysis

#### Respiration preprocessing

In chronic preparations, respiratory activity was inferred from diaphragmatic electromyography (dEMG) in animals implanted with bipolar electrodes in the diaphragm muscle (Rojas-Líbano et al., 2014; Shafford et al., 2006). To isolate the respiratory rhythm, the raw dEMG signal was high-pass filtered (>10 Hz) to emphasize muscle activity, and cardiac artifacts were removed through template subtraction. Specifically, R-peaks were automatically detected from the dEMG. A sliding template was computed by averaging ±10 surrounding cardiac cycles, and this average was subtracted from each R-peak–aligned epoch, suppressing the ECG waveform. The cleaned signal was then rectified and median filtered to obtain a smoothed respiratory envelope. A secondary band-pass filter (0.5–10 Hz) was applied to extract the final respiratory oscillation. This processed signal allowed estimation of instantaneous respiratory frequency and entrainment with olfactory bulb activity across wake, NREM, and REM states.

For acute recordings, inspiratory onsets were detected as the peaks in the second derivative of the calibrated thermistor signal, corresponding to the point of maximal acceleration in temperature drop at the start of inhalation. This marker was used to align LFP traces for respiration-triggered averaging.

#### Sleep scoring

For sleep scoring, a subset of channels was selected. First, all the EEG and EMG channels were loaded into Python. In the case of acute recordings, four distributed LFP channels were also loaded, band-pass filtered (0.5-200 Hz), and down sampled to 1000 Hz and then exported to European Data Format (EDF). Sleep scoring was performed manually using Spike2.v10 software (CED; sleepsco script). Data was obtained during spontaneously occurring W, NREM, and REM sleep. The presence of low voltage fast waves in the EEG, a mixed theta rhythm (4–9 Hz) in the parietal/occipital cortex, and relatively high electromyographic activity were used to identify W. Light and deep NREM sleep were determined, but only epochs of established periods of deep NREM sleep were utilized for analysis. Deep NREM sleep was recognized by the presence of continuous, high-amplitude slow waves (0.5–4 Hz) and sleep spindles (9–15 Hz), combined with reduced EMG activity. REM sleep was identified by the occurrence of low voltage fast parietal waves, a regular theta rhythm in the parietal/occipital cortex, and the absence of EMG Activity except for occasional muscular twitches. A similar scoring process was conducted for the urethane recordings. The occurrence of low voltage fast parietal waves and a regular theta rhythm in the parietal/occipital cortex identified the activated state (ASt). Slow wave state (SWSt) was determined by the presence of continuous high-amplitude slow waves and sleep spindles (as NREM sleep). Finally, the transition (trans) period was identified by the presence of sporadic slow waves with mixed low-voltage fast waves.

#### Coupling analysis

Spectral power, coherence, and phase–amplitude coupling (PAC) were computed using custom scripts in Python. Power spectral density (PSD) and coherence were estimated via multitaper methods (Prerau et al., 2016). PAC between the phase of the respiratory signal and the amplitude of gamma oscillations was quantified using the Modulation Index (MI) as described by *Tort et al., 2010*(Tort et al., 2010).

All signal processing steps, parameters, and code used for generating the figures are publicly available in the project’s GitHub repository (see Data and Code Availability section).

#### Current Source Density (CSD) Analysis

To quantify laminar current flow in the olfactory bulb, we computed one-dimensional current source density (CSD) from local field potentials (LFPs) recorded with linear silicon probes (inter-electrode spacing: 100–200 µm), following the classical method described by Nicholson and Freeman (1975) (Cavelli et al., 2023; Nicholson and Freeman, 1975). Raw LFP signals were first smoothed using a 5-point moving average to reduce spatial noise. The CSD was estimated by applying the second spatial derivative using the three-point formula:

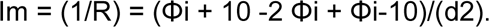

Where Φi is the potential at electrode i, and d = 0.01 mm is the inter-electrode distance. This yields a spatial profile of current sinks (negative values, inward transmembrane current) and sources (positive values, outward current). Boundary channels were excluded from derivative computation to avoid edge artifacts. All computations were performed using custom Python scripts.

### Statistical Analysis

For comparisons involving three or more brain states (e.g., Wake, NREM, REM), we used repeated-measures ANOVA followed by paired t-tests or Wilcoxon signed-rank tests with Bonferroni correction for multiple comparisons. For two-condition analyses, paired t-tests were used when data met normality assumptions; otherwise, Wilcoxon signed-rank tests were applied.

Correlations (e.g., between log delta power and coherence or log MI) were assessed using Pearson’s r. To determine the statistical significance of coupling and coherence values, we generated surrogate datasets by phase-scrambling the signals 200 times and compared observed values to this null distribution.

Statistical analyses were conducted in Python. All reported p-values are two-tailed unless stated otherwise, and significance thresholds were set at *p* < 0.05.

## Data and Code Availability

All custom code and analysis scripts used in this study are available at: [https://github.com/cavelligonca].

Raw data and processed datasets are available from the corresponding author upon reasonable request.

## Acknowledgments

We thank graduate students Juan Pedro Castro and Sofía Niño for their valuable contributions during data collection and analysis. We are also grateful to Dr. Adriano Tort, Dr. Daniel Rojas-Libano, and Dr. Graham Findlay for their insightful comments on the manuscript draft. Special thanks to Dr. Daniel Rojas-Líbano for providing the rat illustration used in several figures, and to Dr. Daniel Olazábal, whose collaboration on an unrelated project indirectly supported the development of this study. Doctoral students S.D., D.G., and M.M. received financial support from the Comisión Académica de Posgrado (CAP), the Agencia Nacional de Investigación e Innovación (ANII), and the Programa de Desarrollo de las Ciencias Básicas (PEDECIBA). This research was supported by multiple grants from the Universidad de la República through the Comisión Sectorial de Investigación Científica (CSIC): CSIC-Grupos (No. 22620220100148UD), CSIC I+D (No. 22520240100405U), CSIC Equipamiento 2024 II (ID#42), and CSIC Equipamiento 2024 III (ID#24).

## Declaration of generative AI and AI-assisted technologies in the writing process

During the preparation of this manuscript the authors used ChatGPT (OpenAI, 2025) to assist in editing language, refining argument structure, and enhancing clarity. The scientific content—including data acquisition, analysis, interpretation, and conclusions—was conceived, validated and approved exclusively by the authors. All authors take full responsibility for the accuracy and integrity of the work.

## Supplementary Material

**Figure S1.**
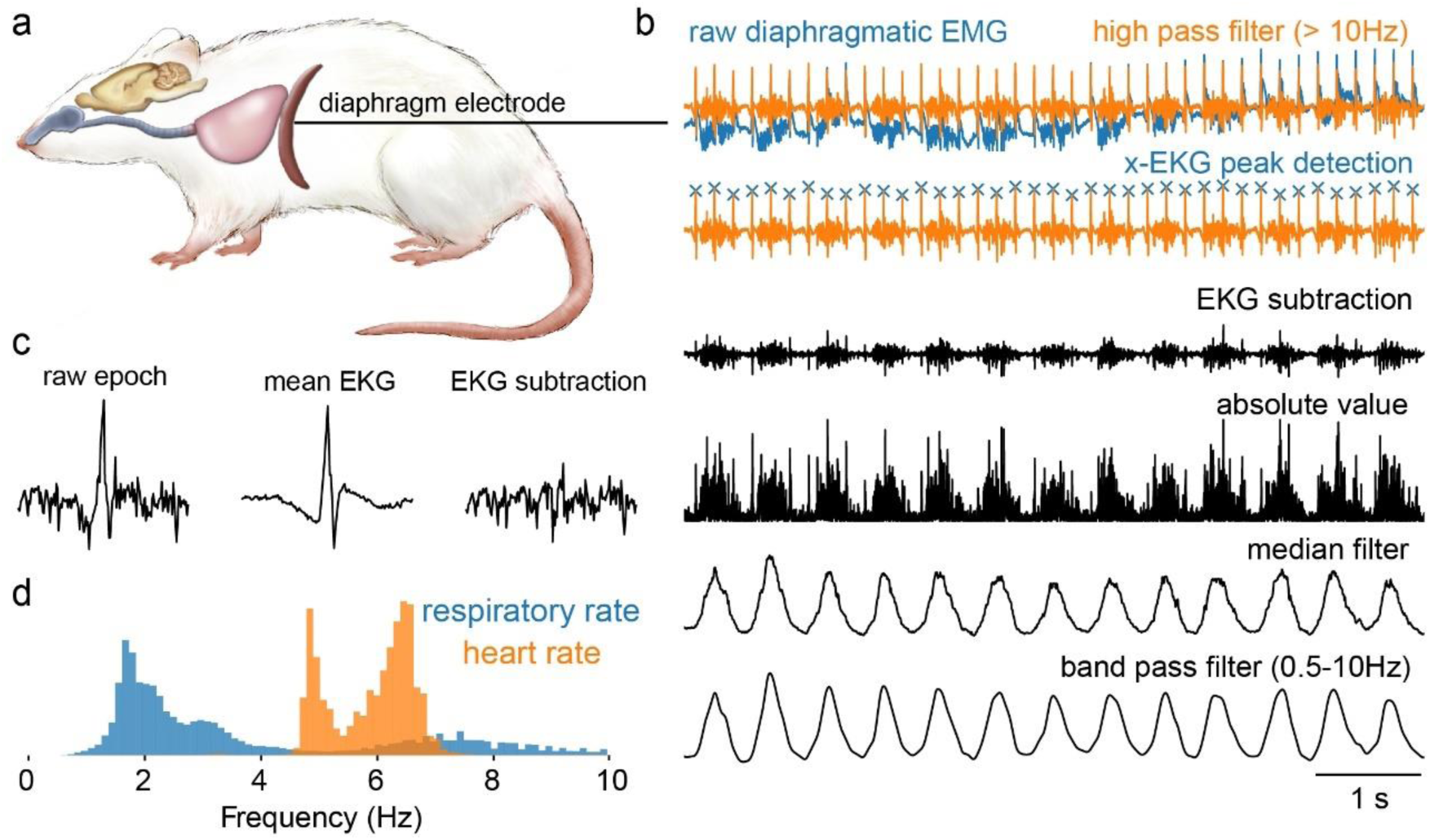
Extraction of respiratory signal from diaphragm EMG with ECG artifact removal. ***a.*** Schematic representation of the experimental setup. A bipolar electrode was implanted in the diaphragm muscle to monitor breathing-related activity (dEMG). ***b.*** From top to bottom: raw diaphragmatic EMG (blue) and the same signal after high-pass filtering above 10 Hz (orange). Detected ECG peaks are marked (×) for subtraction. Post-subtraction trace, absolute value, median filter, and final band-pass filtered signal (0.5–10 Hz), which isolates the respiratory rhythm. ***c.*** Illustration of the ECG subtraction process. Left: an individual raw dEMG epoch. Middle: mean ECG waveform computed across ± 10 detected R-peaks (R-peak moving window). Right: signal after template subtraction. ***d.*** The processed signal allows accurate estimation of the respiratory frequency, clearly separated from the heart rate peak (respiration in blue, heart rate in orange).

**Figure S2.**
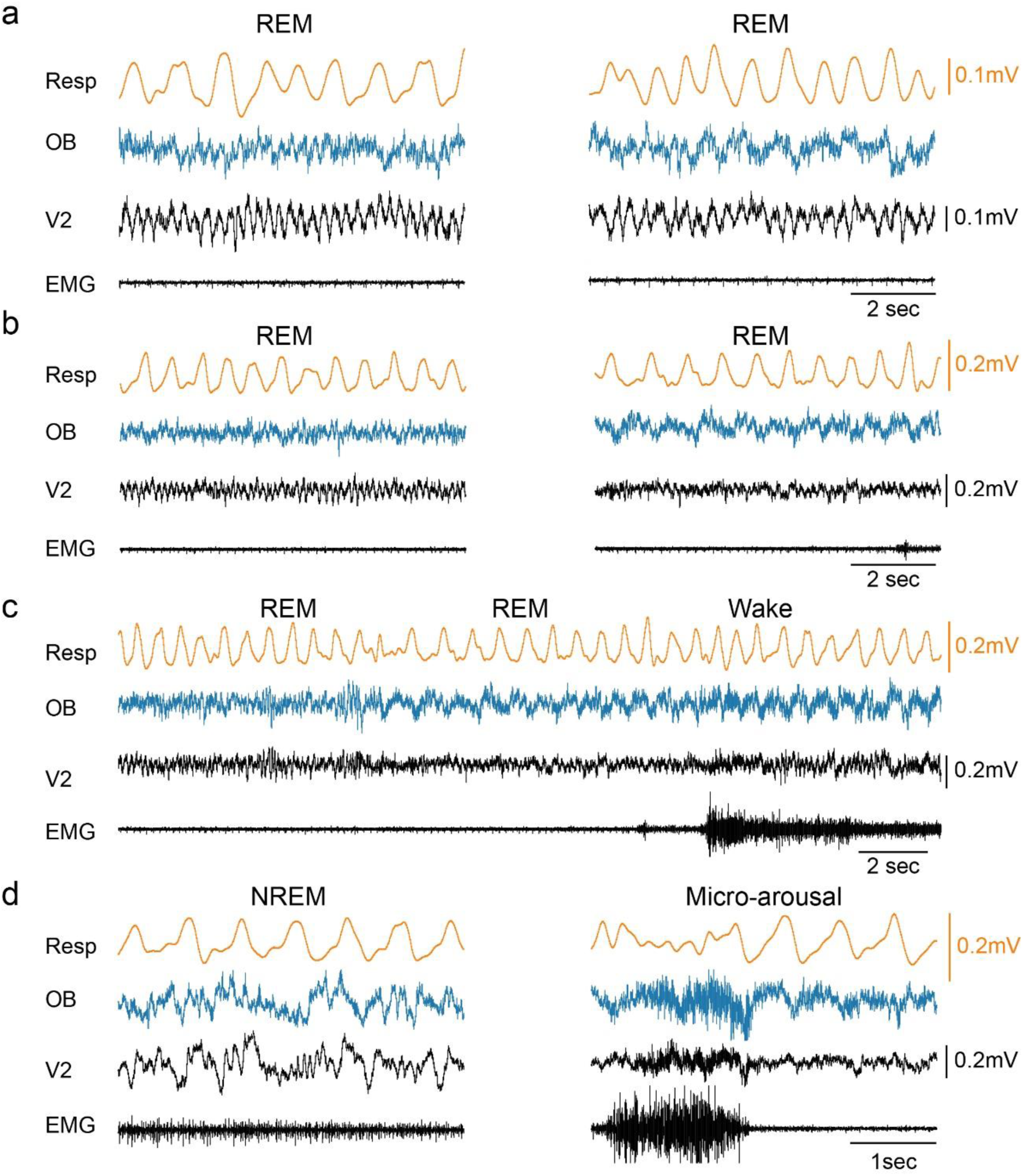
Respiratory-coupled potentials in the OB re-emerge during brief REM sleep periods and microarousals. **a-d.** Example traces of respiration (Resp), olfactory bulb (OB), visual cortex (V2), and EMG during REM and NREM **d.** In each panel, the left trace shows a period with minimal OB respiratory potential. In contrast, the right trace highlights a brief reappearance of the signal, either during short REM sleep periods (a–c) or a microarousal (d). These events illustrate that the respiration-locked potential, while generally absent during NREM and REM sleep, can transiently return during momentary sleep periods.

**Figure S3.**
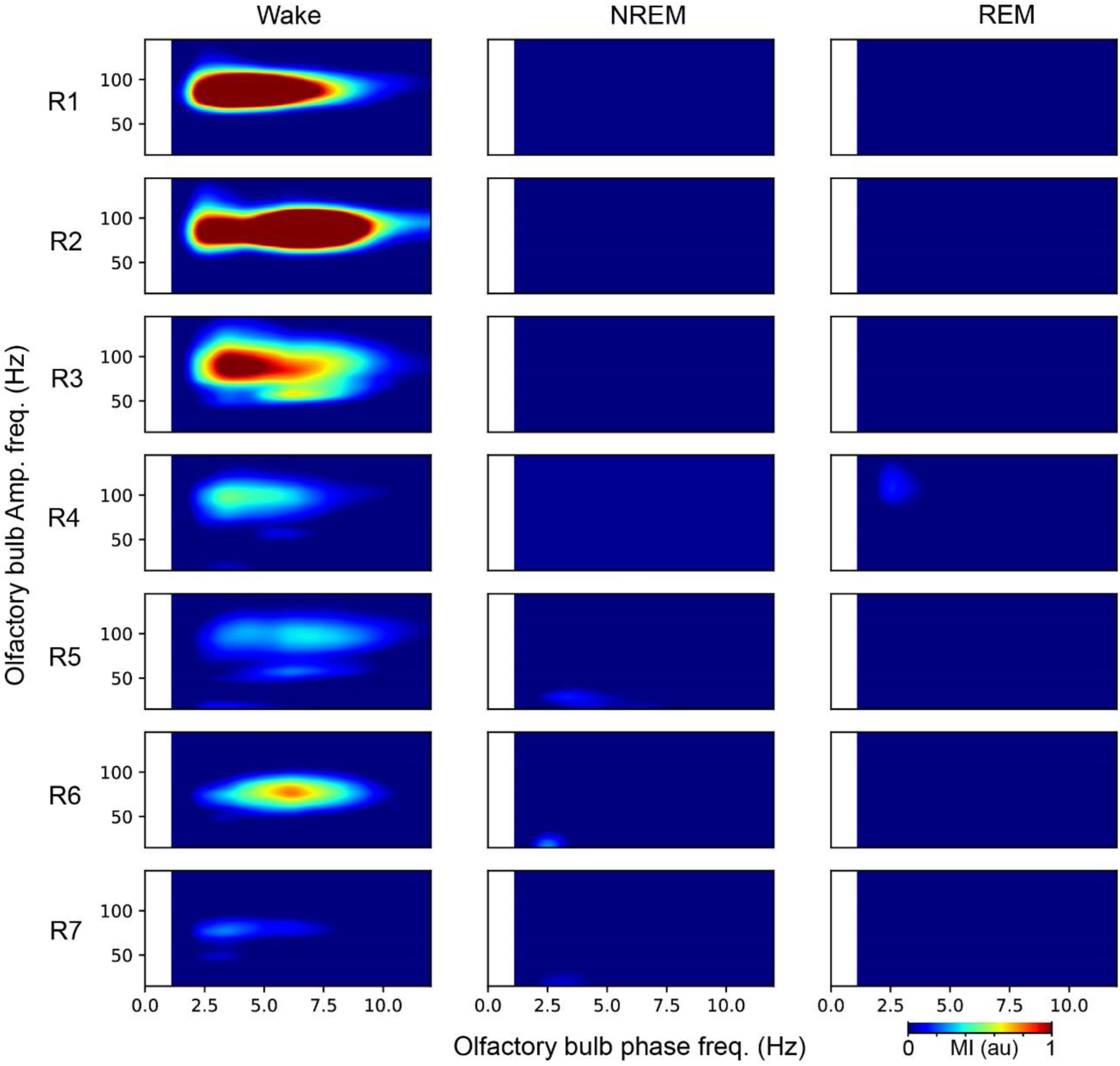
Phase–amplitude coupling (PAC) in the olfactory bulb across sleep-wake states. Comodulograms from seven rats (R1–R7) showing phase-amplitude coupling between low-frequency phase (x-axis) and high-frequency amplitude (y-axis) in the olfactory bulb during Wake, NREM, and REM sleep. During wakefulness, strong PAC is observed between the respiratory-locked potential (∼2-10 Hz) and gamma-band amplitude (∼60–110 Hz), which disappears during both NREM and REM sleep. This suggests a state-dependent decoupling of olfactory bulb circuits from respiratory drive during sleep. Comodulograms were computed using the Modulation Index (MI) method (Tort et al., 2010). au, arbitrary units.

**Figure S4.**
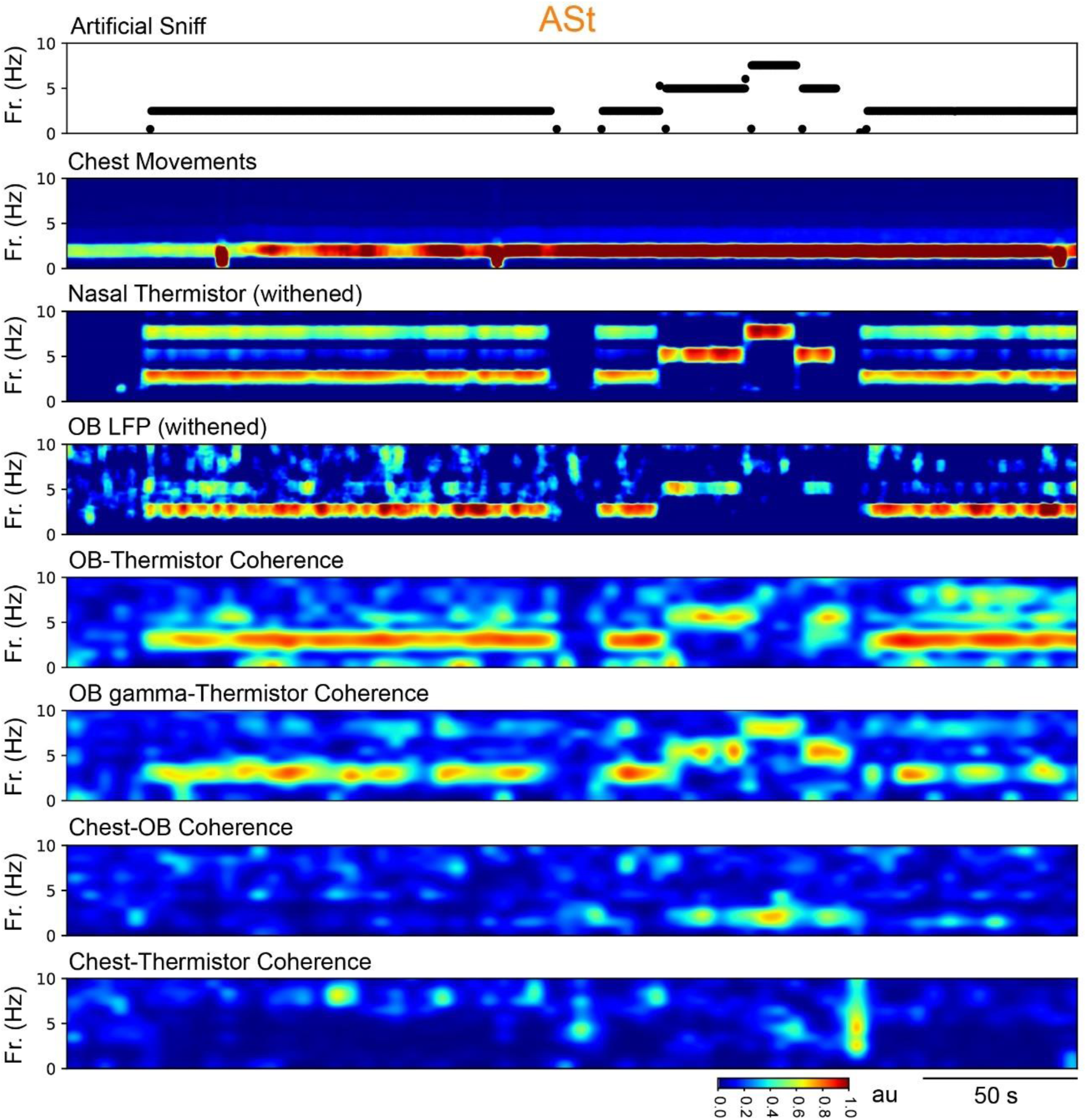
State-dependent olfactory bulb tracking of artificial nasal stimulation during urethane-activated state (ASt). Example recording during ASt with double tracheotomy and artificial sniffing in a chronically implanted rat. The artificial sniff frequency varied throughout the session (top trace), while chest movements and thermistor signals (2nd and 3rd panels from the top) indicate complete respiratory independence between them. The OB LFP (4th panel) and its coherence with the thermistor signal (5th panel) show that stimulus-locked potentials are tightly coupled to artificial sniffing. OB gamma-band activity (6th panel) also follows the artificial sniffing input, suggesting preservation of sensory transmission. In contrast, coherence between chest and OB or thermistor signals (bottom two panels) remains low, confirming effective decoupling of thoracic respiration and nasal air-flow. These findings demonstrate that during ASt, the OB selectively tracks artificial nasal inputs, supporting a gating mechanism that is largely independent of thoracic motor drive. au, arbitrary units.

